# Autonomic regions of the brainstem show a sex-specific inflammatory response to systemic neonatal lipopolysaccharide

**DOI:** 10.1101/2023.06.14.544893

**Authors:** Kateleen E Hedley, Annalisa Cuskelly, Robert J Callister, Jay C Horvat, Deborah M Hodgson, Melissa A Tadros

## Abstract

Early life inflammation has been linked to long-term deficits in the central nervous system in relation to behavioural disorders, but it is now becoming more apparent it can also lead to autonomic dysfunction. The brainstem contains all critical control centres for autonomic homeostasis, so we used the well-established model of neonatal lipopolysaccharide (LPS) exposure to examine the immediate and long-term impacts of systemic inflammation on the autonomic regions of the brainstem. Wistar rats were injected with LPS or saline on postnatal days 3 and 5, with sacrifices made on postnatal days 7 and 90. At both timepoints inflammatory mediators were assessed in the brainstem via RT-qPCR and microglia were characterised by immunofluorescence in the autonomic regions of the brainstem. In the brainstem there was a distinct sex-specific response of all measured inflammatory mediators at both ages, as well as significant neonatal sex differences in inflammatory mediators at baseline. AT both ages, microglial morphology had a significant change to branch length and soma size in a sex-specific manner, which strongly indicate a significant effect of neonatal immune activation. This data not only highlights the strong sex-specific response of neonates to LPS administration, but also the significant impact on the brainstem in adulthood.

## 1. Introduction

The brainstem contains a number of autonomic nervous system (ANS) nuclei crucial for life (Nicholls and Paton, 2009). These control centres, located in the medulla oblongata, modulate the majority of visceral physiology including, but not limited to, heart rate, respiration, gastrointestinal and immune function (Gibbons, 2019). Traditionally, the CNS was considered immune privileged, however, this is no longer the case. Peripheral to central immune communication is now widely recognised, with peripheral inflammation inducing neuroinflammation through a number of well described pathways resulting in the release of inflammatory mediators within the CNS (Shouman and Benarroch, 2021). Peripheral inflammatory mediators interact with the CNS causing neuroinflammation via multiple different pathways, such as through cytokine production and microglia, the resident immune cell in the CNS (Hocker et al., 2017). This disruption to the delicate CNS environment can cause changes to neuronal signalling as well as dysfunction in end organ function. For example, it has been shown that enhanced interleukin 1β (IL-1β) signalling can cause synaptic hyperexcitability in patients with multiple sclerosis (Rossi et al., 2012) and neuroinflammation has been linked to heart failure (Díaz et al., 2020). However, despite the finding that peripheral pro-inflammatory cytokines can stimulate vagus nerve activity (Browning et al., 2017), the impact of inflammation on crucial ANS nuclei in the brainstem remains under studied.

Animal models of early life peripheral inflammation and their effect on the CNS are broad and varied, however, one of the most commonly used is early life exposure to lipopolysaccharide (LPS) (Gilles et al., 1976, Raetz and Whitfield, 2002). LPS is a component of gram-negative bacterial cell walls and activates the innate immune system via toll-like receptor 4 (TLR4) (Nemzek et al., 2008). In contrast to the extensive body of evidence documenting the life-long impact of early life LPS exposure in the brain (Catorce and Gevorkian, 2016), the few studies examining brainstem nuclei only consider the acute impact of LPS (Hedley et al., 2022).. For example, within 24 hrs of LPS exposure there was a significant increase in medullary IL-1β, interleukin 6 (IL-6) and tumour necrosis factor α (TNFα) following LPS exposure (Jafri et al., 2013, Balan et al., 2011) and an increase in the number of microglia in the hypoglossal nucleus (nXII) (Johnson et al., 2016). Further, time- and dose-dependent impairments of nXII output by systemic LPS has been noted in neonatal (P0-6) rats, which corresponded with elevated inflammatory mediators in the medulla oblongata (Morrison et al., 2020). These findings suggest that even low level neuroinflammation can alter the output of brainstem nuclei, confirming that early life systemic LPS exposure can induce acute changes in neuronal function, and subsequently, visceral physiology. Though these studies represent an important foundation, they do not consider any long-term impacts of early life LPS exposure on neuroinflammation in the brainstem. Therefore, in order to fully elucidate the impact of early life events on ANS function, further exploration of the consequences of neonatal inflammation on these key autonomic nuclei is warranted.

Previous studies in our laboratory have shown respiratory deficits in adulthood following systemic neonatal LPS (Sominsky et al., 2013), however the autonomic control centres were not examined and only males were used. In this study, we utilised our well-established model of early life LPS exposure (Cuskelly et al., 2022, Hodgson et al., 2001, Tadros et al., 2018) to examine the immediate and long-term impacts of systemic inflammation on the autonomic regions of the brainstem. We hypothesised that neonatal immune activation via LPS exposure would modify the inflammatory profile in the brainstem, in both neonates and adults. We assessed a number of inflammatory mediators via RT-qPCR and examined the morphological characteristics of microglia to determine activation status, as the immune mediators of the CNS. Our findings demonstrate strong sex-specific responses of all measured inflammatory mediators at both ages, as well as significant neonatal sex differences in inflammatory mediators at baseline. The morphological modifications, such as changes to branch length and soma size of microglia, strongly indicate a significant effect of neonatal immune activation on at both neonatal and adult ages. This data not only highlights the strong sex-specific response of neonates to LPS administration, but also the significant impact on the brainstem in adulthood

## 2. Materials and Methods

### 2.1. Animals

All experiments were performed in accordance with the National Health and Medical Research Council Australian Code of Practice and as approved by the University of Newcastle Animal Care and Ethics Committee (approval number A-2018-814). Naïve male and female Wistar rats were acquired from the University of Newcastle’s Animal House aged between 10-12 weeks. The ratio used for breeding was 1 male to 4 females, with the rat pups born to these dams used as the experimental animals (n = 91; females = 47, males = 44). All animals were maintained in a temperature (22 °C ± 2) and humidity-controlled environment on a 12hr light/dark cycle with food (standard rat chow) and water available ad libitum for the duration of the study.

### 2.2. Neonatal immune activation model

The laboratory model of neonatal immune activation used in this study has been previously published (Cuskelly et al., 2022). Briefly, on the day of birth, assigned postnatal day 1 (P1), all pups within a litter were randomly allocated to receive either lipopolysaccharide (LPS) or saline (SAL). Pups were given an intraperitoneal injection (i.p.) of either 0.05 mg/kg of LPS (pyrogen free saline and *Salmonella entricia*, serotype Enteritidis) or saline (equivolume) on P3 and again on P5 then left undisturbed with their dams until the selected timepoints of P7 or P90, to assess both the acute and long-term impacts of neonatal LPS-exposure. Neonatal animals were sacrificed via guillotine and tissue obtained, P90 animals were anaesthetised via isoflurane inhalation and then sacrificed via guillotine, with tissue then collected. For both timepoints and tissue types the experimental groups were as follows; male LPS, male SAL, female LPS, female SAL.

### 2.3. RNA extraction and RT-qPCR for inflammatory markers

Brainstems and spleens (n=6-8 per group) were removed following euthanasia and immediately snap-frozen to -80°C. The medulla oblongata was then separated from the whole brainstem and homogenised, and the whole spleen was homogenised with a portion of the resulting homogenate (10-20mg) taken for further processing. Total cellular RNA was extracted using the RNeasy® Mini Kit (Qiagen, Hilden, Germany), following manufactures instructions. RNA concentration and quality were measured with a NanoDrop 1000 Spectrophotometer (ThermoFisher Scientific, USA).

Any contaminating genomic DNA remaining in the RNA samples was digested using DNase I (Invitrogen, Scoresby, Australia). Reverse transcription was then performed using Superscript III (Invitrogen, Scoresby, Australia), according to manufacturer’s instructions.

Briefly, 30 - 200ng of total RNA, 1µl of oligo(dT)_18_ (Meridian Bioscience, Ohio, USA), 1µl of random hexar (Meridian Bioscience, Ohio, USA), 1µl of 10µM dNTP (Meridian Bioscience, Ohio, USA), and molecular grade water to 13µl, were mixed and heated for 5 min at 65°C in a Thermal Cycler (Eppendorf, Germany). Next, 4µl of 5x first-strand buffer, 1µl of 0.1M DTT, 1µl RNaseOUT (40 U/µl) (Meridian Bioscience, Ohio, USA) and 1 µl SuperScript III RT (200 U/µl) were added and the mixture was incubated for 60 min at 50°C, followed by 70°C for 15 min. Reverse transcription without Superscript III was also performed to assess genomic DNA contamination.

All qPCR primers (Table 1) were designed with Ensembl using the standard primer design criteria. The primer pairs were then put through NCBI primer BLAST to ensure primer specificity. A total volume of 12.5µl was used containing: 6.25µl 2x SensiFAST SYBR (Meridian Bioscience, Ohio, USA), 10µM each of forward and reverse primers, 1 - 2.5ng cDNA and molecular grade water to 12.5µl. After an initial 10 min 95°C enzyme activation step, 40 cycles of 95°C for 30 s (step 1) followed by 60°C for 30 s (step 2) were completed. Melt curves were generated to confirm the presence of a single PCR product. Primers were deemed specific if a single amplified product of appropriate size was detected by melt curve analysis. Reactions were performed on a 7500 Real Time PCR System (Applied Biosystems, USA) and analysed using the Applied Biosystems 7500 Software (version 2.3). For each primer the samples were run on a 96-well plate in triplicate, including a negative water control on each plate. Delta Ct (ΔCt, threshold cycle) was determined for each gene relative to the housekeeping genes β-Actin and 18S, and the well-established ΔΔCt method (Livak and Schmittgen, 2001) was employed to enable comparisons between the groups.

**Table 1.**
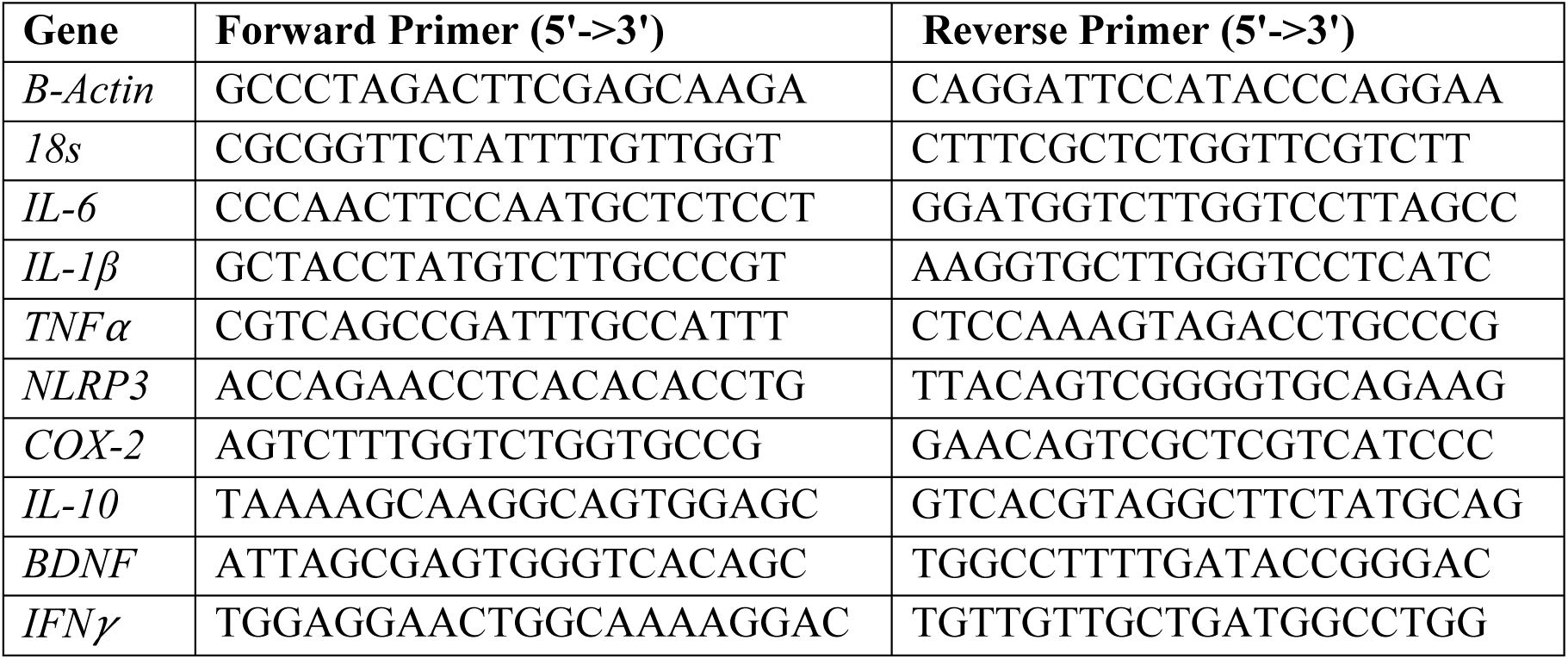
Forward and reverse primer sequences.

### 2.4. Immunofluorescent labelling of microglia

Due to unexpected litter sizes, a small surplus of animals were available, therefore brainstems (n=3-4 per group) were dissected following euthanasia and immersion-fixed in 4% paraformaldehyde (PFA) in phosphate buffer for 24hrs then washed into phosphate buffered saline (PBS). Brainstems were then washed with 3 x 15min 80% ethanol (EtOH), 3 x 15min dimethyl sulfoxide (DMSO) and then 3 x 15min 100% EtOH. The brainstems were then placed in a cell culture dish containing melted 1000MW polyethylene glycol (PEG) in a 46°C vacuum oven (Thermoline, Wetherill Park, NSW) until the tissue had sunk (approximately 2 hours). The tissue was then placed in a cryostat mould and embedded in 1450MW PEG. Once hardened, the tissue block was sectioned at 30μm on a rotary microtome. Every one in four sections from bregma level -12.80mm to -14.60mm were taken and placed in PBS. The sections were then blocked in 10% normal donkey serum (NDS, Jackson ImmunoResearch) for 30min. For labelling, sections were incubated at room temperature overnight in a solution of antibody diluent (0.3M NaCl, 7.5mM Na_2_HPO_4_, 2.5mM NaH_2_PO_4_, 0.3% TritonX-1000, 0.05% Na Azide), 10% NDS and ionized calcium binding adaptor molecule 1 (Iba1; rabbit; 1:250; Wako; 019-19741). After the primary incubation, sections were rinsed in 3 x 15min PBS and incubated for 2hrs at room temperature with secondary antibodies Alexa594-donkey-anti-rabbit (1:50; Abcam: ab150076) in antibody diluent. Sections were mounted on slides in buffered glycerol and cover slipped. Images were acquired using an Olympus BX50 microscope equipped with a mercury burner and an Olympus DP72 camera. A brain atlas was used to identifying to specific nuclei and they were cut from the images using a cutting tool in ImageJ. Images were processed offline using a custom MATLAB script to analyse microglia number and morphology (Abdolhoseini et al., 2019, Abdolhoseini et al., 2016, Kluge et al., 2017).

### 2.5. Statistics

For RT-qPCR, statistical differences were determined using a multivariate analysis of variance (ANOVA) using SPSS. Fisher’s least significant difference (LSD) was used for post-hoc comparisons, with significance set at p < 0.05. Due to small sample size, Cohen’s D was the most appropriate statical comparison for microglia immunofluorescence, with effect sizes (d_s_) being defined as large >0.8, moderate >0.5 and small >0.2. All graphs presented use untransformed means and standard errors of means (SEM).

## 3. Results

This study focussed on two specific timepoints: two days after LPS exposure, at P7, in order to examine the acute impact of neonatal exposure; and during adulthood, at P90, to determine any long-term alterations following neonatal exposure to LPS. Utilising spleen and brainstem allowed a comparison between peripheral and central responses to neonatal LPS.

### 3.1 IL-6 altered in both the spleen and brainstem after neonatal LPS exposure

We first examined interleukin 6 (IL-6), a pro-inflammatory cytokine, as it is readily released and is involved in the acute phase of the immune response to LPS (Tanaka et al., 2014). We investigated the levels of IL-6 mRNA in the spleen and brainstem at both timepoints in this study as an indicator of peripheral and central inflammation, respectively. At P7, treatment × sex analyses of the spleen and brainstem showed sex-specific effects of LPS exposure in both tissues. In the spleen, LPS exposed males had a significant increase in IL-6 expression compared to saline-treated males, as shown in Fig. 1A (post-hoc for males F _(1,26)_ = 14.629, p < 0.001). In contrast, the brainstems of LPS exposed males had a significant decrease in IL-6 expression compared to saline males, as shown in Fig. 1D (post-hoc for male F _(1,26)_ = 14.961, p < 0.001). Surprisingly, there was no impact of LPS-exposure on female spleens or brainstems at P7 (post-hoc for female spleens F _(1,26)_ = 2.441, p = 0.130, Fig. 1A; post-hoc for female brainstems F _(1,26)_ = 3.256, p = 0.083, Fig. 1D). Interestingly, in the brainstem a sex x treatment effect was observed, revealing baseline differences between the males and females treated with saline (post-hoc for saline F _(1,26)_ = 13.147, p = 0.001, Fig. 1D). These results highlight the differences between central and peripheral immune modulation, and indicate sex differences exist in the naïve neonatal brainstem as well as in the acute response to LPS exposure.

**Figure 1.**
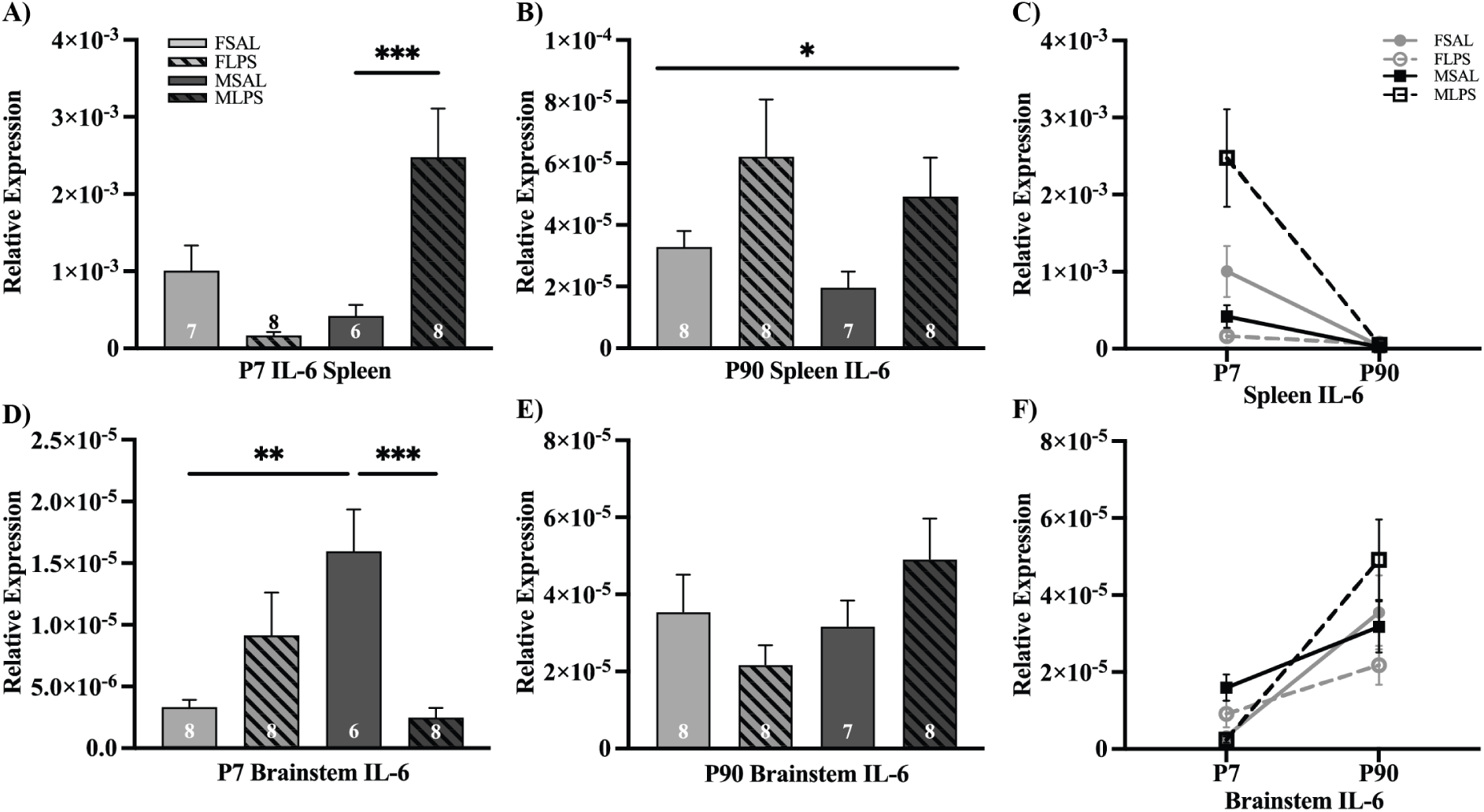
Relative expression of IL-6 in spleen and brainstem homogenates following neonatal LPS. Graphs show IL-6 expression (mean ± SEM) for saline-exposed (open bars) and LPS-exposed (hatched bars) females (light) and males (dark grey), with total number of samples shown in each bar. A) IL-6 expression in the spleen at P7, as a treatment by sex effect (males ***p< 0.001). B) IL-6 expression in the spleen at P90, treatment effect (males *p = 0.022). D) IL-6 expression in the brainstem at P7, as a treatment by sex effect (males ***p< 0.001) and a sex by treatment effect (SAL **p= 0.001). E) IL-6 expression in the brainstem at P90, no significant effects seen. C) and F) show the age-based changes of IL-6 expression in the LPS model in both the spleen and brainstem, respectively.

In adulthood (P90), the spleen of LPS exposed animals showed a significant main effect of treatment on IL-6 expression, as shown in Fig. 1B (Treatment effect: F _(1,27)_ = 5.937, p = 0.022), that was found to be independent of sex. These findings demonstrate a persistently elevated IL-6 in the periphery following neonatal LPS exposure which remains high throughout the animal’s life span. In contrast, in the brainstem there was no significant difference between LPS or SAL groups of either sex, or between the sexes in either group (Fig. 1E), with the expression of IL-6 in the adult brainstem unaffected by neonatal LPS. The levels of IL-6 were compared between the two timepoints for all groups, in order to assess the changes in IL-6 expression with and without LPS (Fig. 1C, F). Overall, in the spleen, there was a reduction in IL-6 expression, with males exposed to neonatal LPS showing a sharper decline in expression levels. In contrast, there was a difference between the two timepoints in IL-6 expression in the brainstem, with males exposed to neonatal LPS again showing the sharpest change in expression levels. Taken together, these results indicate that peripheral and central tissues display alternate expression profiles of IL-6, and that these levels are altered in a tissue-specific manner following neonatal exposure to LPS.

### 3.2 Inflammatory mediators measured in the brainstem show sex-based differences at neonatal and adult timepoints following LPS exposure

#### 3.2.1 NLRP3, IL-1β and TNFα

The NLR family pyrin domain containing 3 (NLRP3) is a major component of the inflammasome and triggers the immune response. Once triggered, NLRP3 activates caspase-1 which then cleaves pro-interleukin 1β into its mature form, the highly pro-inflammatory IL-1β (Latz et al., 2013). Tumour necrosis factor α (TNFα), while also acting as a pro-inflammatory cytokine, has been shown to modulate caspase-1 activity and subsequently alter IL-1β production (Álvarez and Muñoz-Fernández, 2013). These three mediators act in concert as part of the initial inflammatory response. At P7, treatment x sex analyses of NLRP3 and IL-1β expression showed that both inflammatory markers were altered by both sex and treatment. LPS-exposed males had a significant decrease in NLRP3 and IL-1β expression compared to saline-exposed males, as shown in Fig. 2A (post-hoc for males, NLRP3; F _(1,26)_ = 5.208, p = 0.031, IL-1β; F _(1,26)_ = 8.229, p = 0.008). In contrast to the decreases observed in males, LPS-exposed females exhibit a significant increase in both NLRP3 and IL-1β expression (post-hoc for females, NLRP3; F _(1,26)_ = 6.656, p = 0.016, IL-1β; F _(1,26)_ = 4.971, p = 0.035). There was also a sex x treatment effect in both NLRP3 and IL-1β, showing significantly higher baseline levels in saline-treated males relative to saline-treated females (post-hoc for saline, NLRP3; F _(1,26)_ = 7.701, p = 0.010, IL-1β; F _(1,26)_ = 11.624, p = 0.002). LPS exposure also altered TNFα levels, however there were no sex-specific differences as both males and females exposed to LPS showed decreased levels of TNFα compared to their saline-treated counterparts (post hoc for males; F _(1,26)_ = 4.445, p = 0.045, females; F _(1,26)_ = 6.632, p = 0.016). These results demonstrate the NLRP3 inflammasome pathway is rapidly affected by neonatal exposure to LPS.

**Figure 2.**
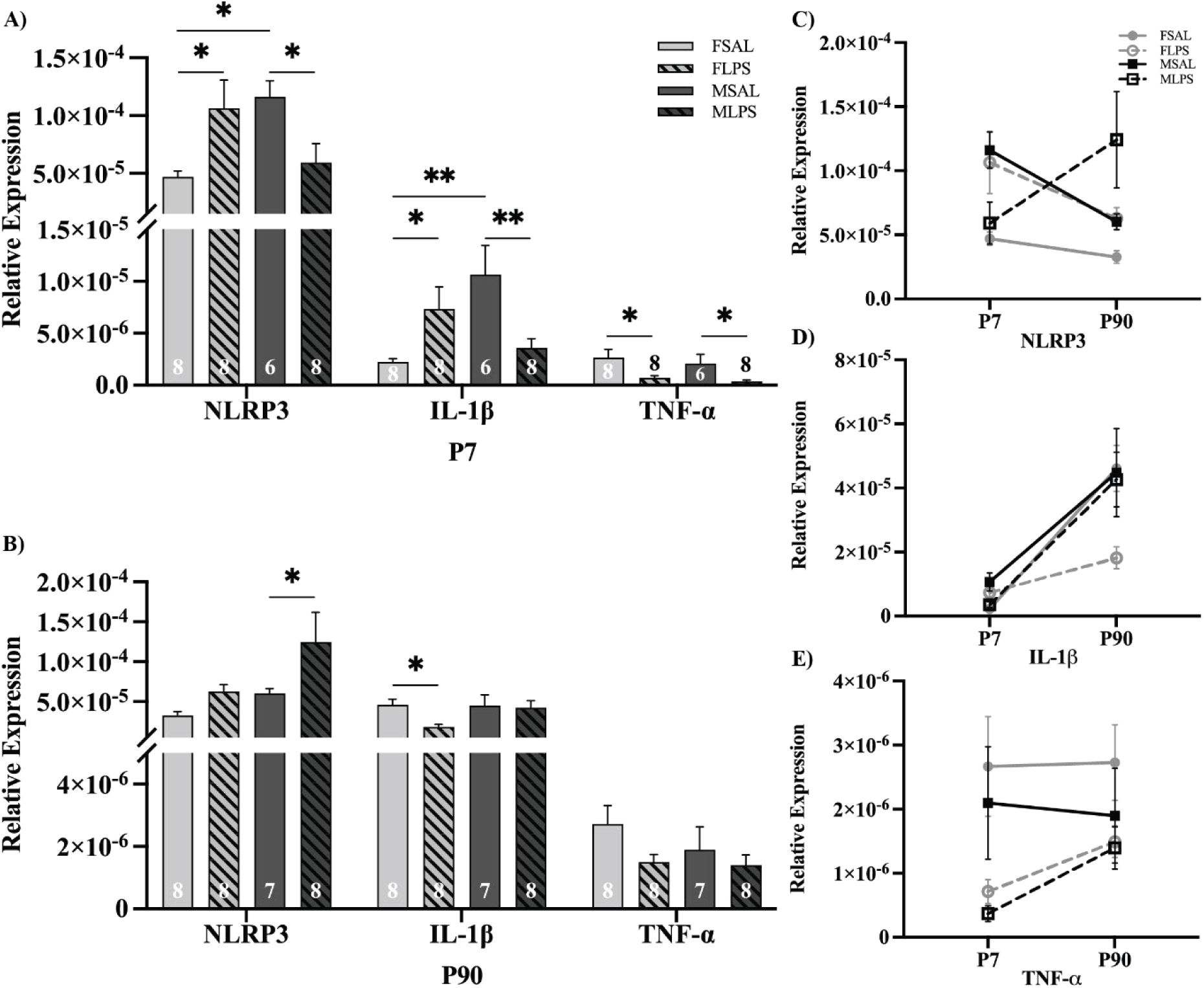
Relative expression of NLRP3, IL-1β and TNFα in the brainstem. Graphs show NLRP3, IL-1β and TNFα expression levels (mean ± SEM) for saline-exposed (open bars) and LPS-exposed (hatched bars) females (light) and males (dark grey), with total number of samples shown in each bar. A) NLRP3, IL-1β and TNFα expression at P7, as a treatment by sex effect (*p<0.05, **p< 0.01). B) NLRP3, IL-1β and TNFα expression at P90, as a treatment by sex effect (*p < 0.05). C-E) show the age-based changes of NLRP3, IL-1β and TNFα expression in the LPS model.

At P90 (Fig. 2B), treatment x sex analyses also revealed sex-specific effects of treatment in ways that differed from the affects observed at P7. Neonatal LPS exposure did not alter NLPR3 levels in adult females, but NLRP3 was elevated in LPS-exposed male adults (post-hoc for males F _(1,27)_ = 4.778, p = 0.038). In opposition to NLRP3, IL-1β levels were unchanged in adult males, but decreased in LPS-exposed female adults (post-hoc for females F _(1,27)_ = 5.447, p = 0.027). TNFα expression did not differ between any of the adult groups examined. By comparing the two ages, the long-term impact of neonatal LPS exposure was revealed. Interestingly, NLRP3 (Fig. 2C) was observed to decrease in all groups, except for males exposed to neonatal LPS, which demonstrated an increase in adulthood. Generally speaking, IL-1Β increased in the brainstem from P7 to P90, however the increase was dampened by LPS exposure in females (Fig 2D). Expression levels of TNFα remained rather constant in saline-treated animals, but showed a relative increase in adulthood in LPS-treated animals (Fig. 2E). The disruption of multiple elements in a key inflammasome pathway clearly highlights the compounding effect of neonatal LPS exposure on both the immediate and long-term regulation of inflammation within the brainstem.

#### 3.2.2 COX-2 and IL-10

To obtain a thorough assessment of multiple inflammatory pathways, we next assessed the levels of interleukin 10 (IL-10) and cyclooxygenase 2 (COX-2) in brainstems from rats exposed to neonatal LPS. IL-10 is a strong anti-inflammatry cytokine that can modulate the expression of pro-inflammatory cytokines, as well as modulating the production of COX-2 (Berg et al., 2001). COX-2 is an enzyme responsible for the production of prostaglandins, which are heavily implicated in a number of physiological responses to inflammation. (Chen, 2010). At P7, treatment x sex anaylsis of COX-2 expression revealed sex-specific effects of treatment. LPS-exposed males demonstrated a significant decrease in expression (Fig. 3A; post-hoc for males F _(1,26)_ = 5.392, p = 0.028), however LPS-exposed females showed no difference to the saline-treated females (post-hoc for females F _(1,26)_ = 3.589, p = 0.069).

**Figure 3.**
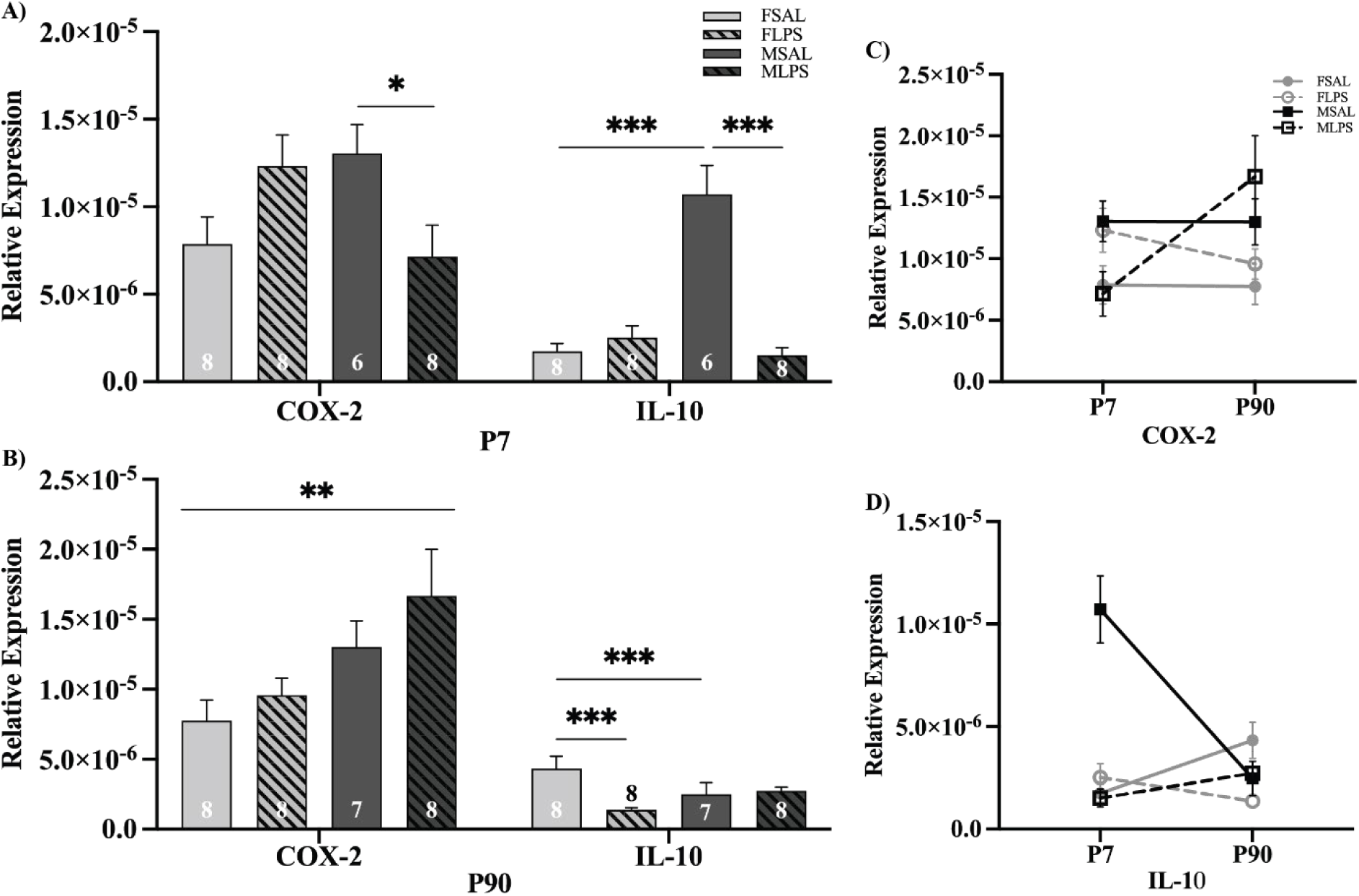
Relative expression of COX-2 and IL-10 inflammatory markers. Graphs show COX-2 and IL10 expression (mean ± SEM) for saline-exposed (open bars) and LPS-exposed (hatched bars) females (light) and males (dark grey), with total number of samples shown in each bar. A) COX-2 and IL-10 expression at P7, as a treatment by sex and sex by treatment effect (*p < 0.05, ***p < 0.001). B) COX-2 and IL-10 expression at P90, as a treatment by sex and sex by treatment effect (*p < 0.05, ***p < 0.001). C, D) show the age-based changes of COX-2 and IL-10 expression in the LPS model.

Expression of IL-10 at P7 showed both sex x treatment and treatment x sex effects, showing sex-specific responses to LPS as well as baseline differenecs between the sexes. Similar to COX-2, LPS-exposed males also showed a decrease in IL-10 expression (Fig. 3A; post-hoc for males F _(1,26)_ = 59.571, p < 0.001), with no impact of LPS observed in females (post-hoc for females F _(1,26)_ = 0.504, p = 0.484). Interestingly, in addition to modulation following LPS exposure, a difference in baseline levels of IL-10 were observed in saline-treated animals, with male brainstems displaying significantly higher expression compared to females (post-hoc for saline, F _(1,26)_ = 56.698, p < 0.001).

Neonatal LPS exposure also altered the expression of COX-2 and IL-10 into adulthood. At the P90 timepoint in COX-2, there was a significant main effect of sex (sex effect: F _(1,27)_ = 8.187, p = 0.008), that was found to be independent of treatment (Fig. 3B), suggesting that males and females have distinct profiles of COX-2 expression. IL-10 expression levels were also altered in adulthood, however, the direction of the impact of neonatal LPS was opposite to P7. At baseline, saline-treated females showed higher levels of IL-10 than males (post-hoc for saline, F _(1,27)_ = 4.481, p = 0.044). Neonatal LPS exposure resulted in a decreased expression of IL-10 in adult females (post-hoc for female, F _(1,27)_ = 12.140, p = 0.002), but no change in adult males exposed to LPS (post-hoc for males, F _(1,27)_ = 0.92, p = 0.764).

A direct comparison of the levels of COX-2 expression between P7 and P90 (Fig. 3C) revealed very little change over time in saline-treated rats, although males displayed a consistently higher expression than females. In the LPS-exposed animals, both males and females showed a greater deviation from saline-treated at P7 compared to P90, suggesting an attempt to correct the alterations resulting from early life exposure by more closely aligning to their saline-treated counterparts at P90. The alteration of IL-10 expression was more complex, with saline-treated females decreasing and saline-treated males increasing with age. LPS-exposure further alter this, with opposing changes observed in LPS-exposed animals compared to saline-treated animals. Clearly, neonatal exposure to LPS has a sex-specific impact on the brainstem expression of IL-10 and COX-2, however the contribution of these two inflammatory mediators is complex and less heavily impacted than other inflammatory pathways.

#### 3.2.3 BDNF and IFNγ

Brain derived neurotrophic factor (BDNF) is a key component in brain plasticity and neuronal development, growth and survival, and is implicated in establishing functional neuronal networks during development (Bathina and Das, 2015). Interferon γ (IFNγ) is a pro-inflammatory cytokine that can promote the secretion of BDNF (Abd-El-Basset et al., 2020). At P7, treatment x sex analyses of BDNF and IFNγ expression showed that both inflammatory markers have sex-specific effects of treatment. LPS-exposed males had a significant decrease in BDNF and IFNγ expression compared to saline males, as shown in Fig. 4A (post-hoc for males BDNF; F _(1,26)_ = 48.067, p < 0.001, IFNγ; F _(1,26)_ = 28.557, p < 0.001). In contrast, there was no difference between in LPS-exposed and saline-treated females for either BDNF or IFNγ (post-hoc for females BDNF; F _(1,26)_ = 0.419, p = 0.523, IFNγ; F _(1,26)_ = 0.020, p = 0.888). BDNF and IFNγ expression also demonstrated sex x treatment effects, with saline-treated males showing higher expression than females for both mediators, highlighting baseline differences exist between the sexes (post-hoc for saline BDNF; F _(1,26)_ = 21.346, p < 0.001, IFNγ; F _(1,26)_ = 19.164, p < 0.001).

**Figure 4.**
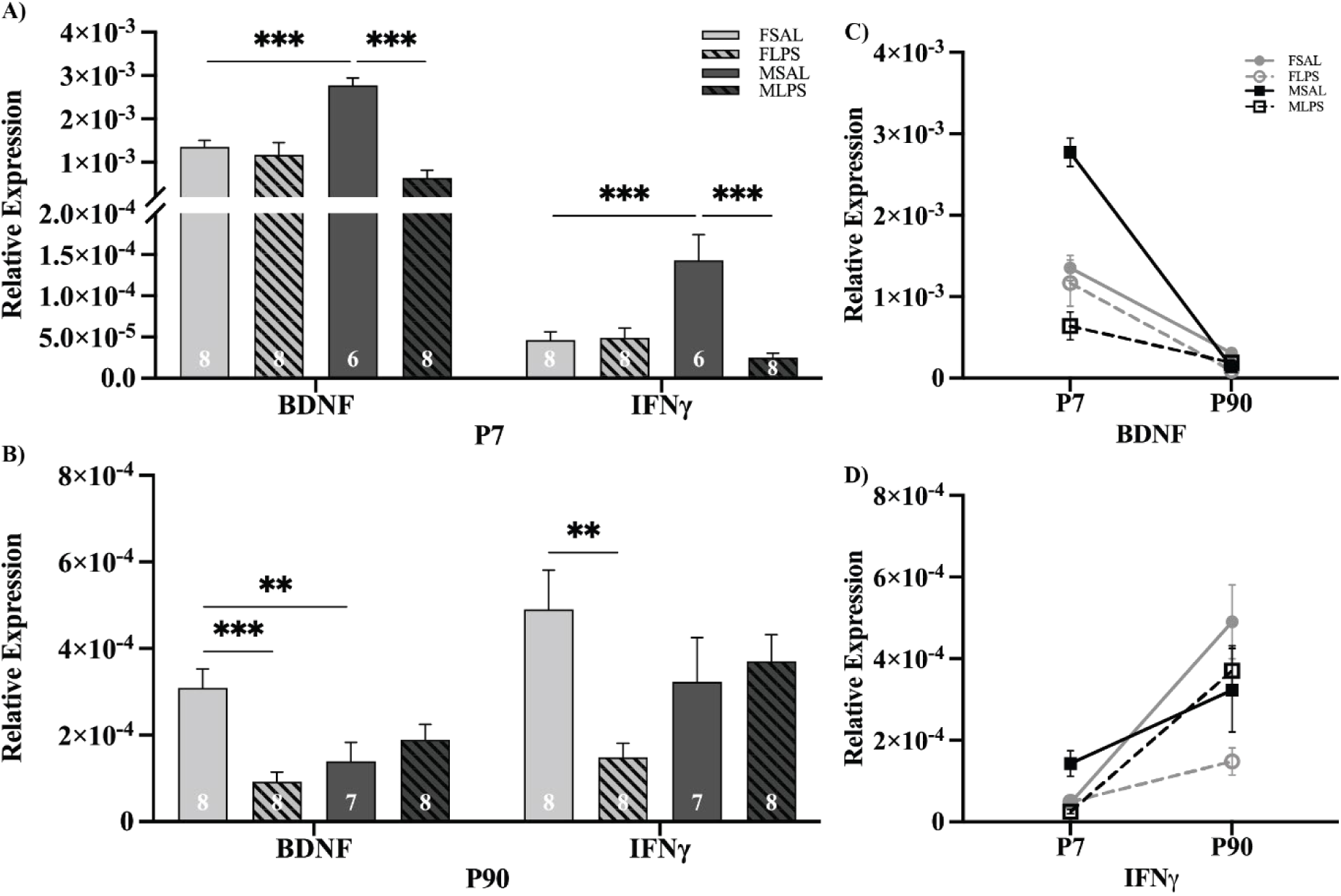
Relative expression of BDNF and IFNγ. Graphs show BDNF and IFNγ expression (mean ± SEM) for saline-exposed (open bars) and LPS-exposed (hatched bars) females (light) and males (dark grey), with total number of samples shown in each bar. A) BDNF and IFNγ expression at P7, as a treatment by sex and sex by treatment effect (***p < 0.001). B) BDNF and IFNγ expression at P90, as a treatment by sex and sex by treatment effect (**p < 0.01, ***p < 0.001). C, D) show the age-based changes of BDNF and IFNγ expression in the LPS model.

In adulthood, treatment x sex analyses of BDNF and IFNγ expression showed that both inflammatory markers exhibit sex-specific effects on treatment. Contrary to P7, the effect was observed in LPS-exposed females, shown in Fig. 4D (post-hoc for females BDNF; F _(1,27)_ = 17.703, p < 0.001, IFNγ; F _(1,27)_ = 10.832, p = 0.003), with no impact of neonatal LPS-exposure on P90 males. The sex x treatment analysis of BDNF expression at P90 again revealed sex differences in saline-treated animals, but the differences were reversed, with females showing higher brainstem expression than males (post-hoc for saline; F _(1,27)_ = 10.145, p = 0.004). Comparing P7 and P90 timepoint of BDNF and IFNγ demonstrated all age-related changes occurred in similar directions, with BDNF decreasing and IFNγ increasing over time. Although LPS exposure altered levels of these mediators, the overall direction of change into adulthood remained similar between saline- and LPS-treated animals.

### 3.3 Microglia are significantly altered in adulthood follow neonatal LPS

Following the revelation of altered expression of inflammatory mediators after neonatal LPS exposure, pilot data for microglial activity was assessed to attempt to elucidate an underlying mechanism. We examined three specific brainstem nuclei: the dorsal motor nucleus of the vagus (DMX) and the solitary tract of the nucleus (NTS) to represent critical centres for autonomic control of viscera and the hypoglossal nucleus (nXII) as a representative somatic control region. In order to best determine the impact of LPS exposure on a small sample of brainstems, Cohen’s d_s_ statistics and consideration of the resulting effect sizes were the most appropriate statistics. The largest effects were seen in the DMX (Fig 5 and 6), however the NTS and nXII followed similar trends (Supp Fig 1-4). DMX data is presented in the main portion of this article and NTS and nXII data can be found in the supplementary data due to the similar trends found for each nuclei.

**Figure 5.**
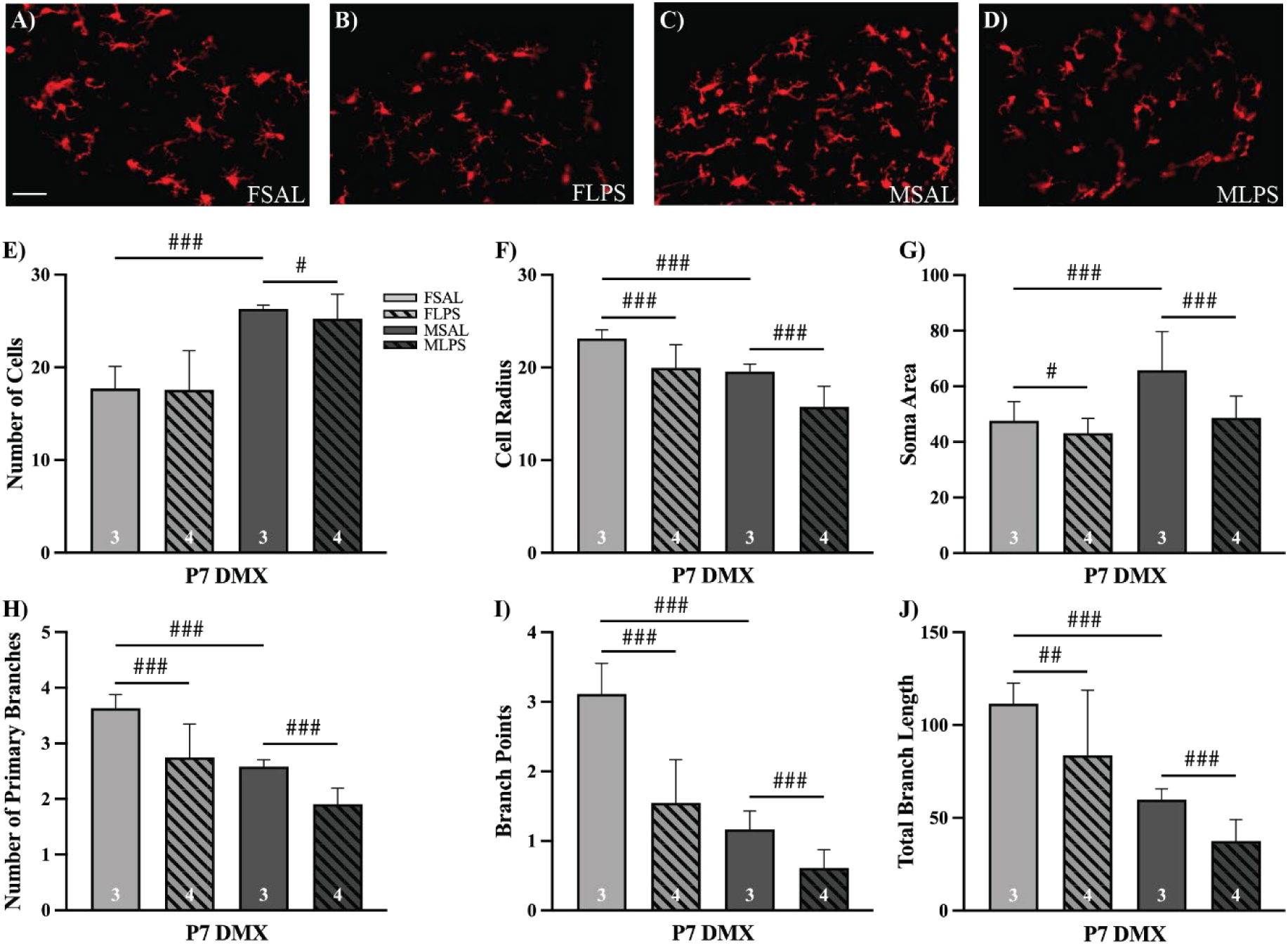
Immunofluorescence of microglia within the dorsal motor nucleus at P7. A-D) Representative images of Iba1 fluorescent labelling for microglia in all four groups. Scale bar 100µm. Bar graphs showing microglia morphological characteristics (mean ± SEM) for Saline-exposed (open bars) and LPS-exposed (hatched bars) females (light) and males (dark grey), with total number of samples in the bottom of each bar. All image analysis was completed using a custom MATLAB script and quantified the following characteristics (all units in pixels) E) Number of cells, F) Cell radius, G) Soma Area, H) Number of primary branches, I) Branch points or secondary branches, J) Total branch length. Effect size statistics were run and can be seen in Table 2, large = ###, moderate = ##, small = #.

**Figure 6.**
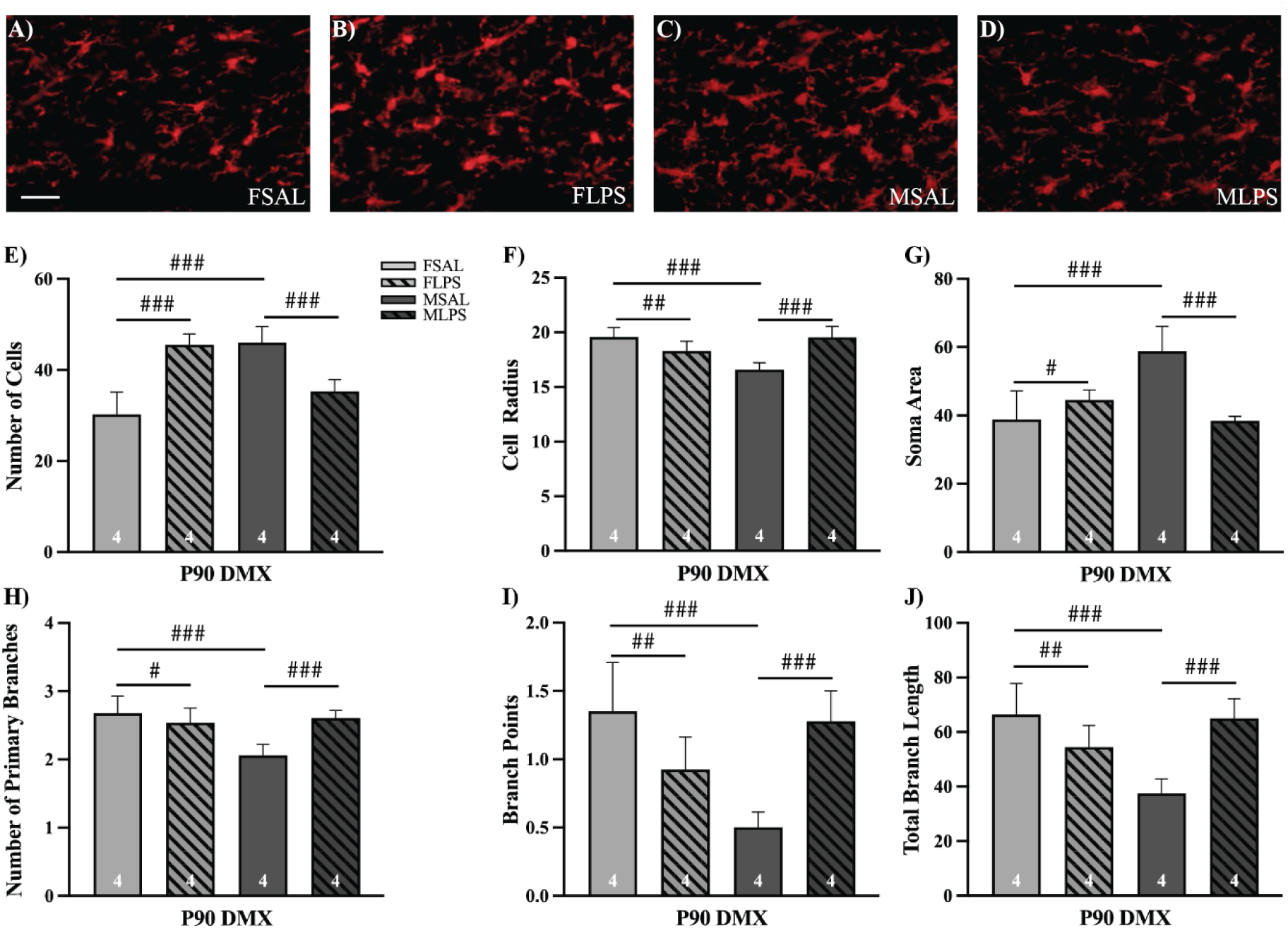
Immunofluorescence of microglia within the dorsal motor nucleus at P90. A-D) Representative images of Iba1 fluorescent labelling for microglia in all four groups. Scale bar 100µm. Bar graphs showing microglia morphological characteristics (mean ± SEM) for Saline-exposed (open bars) and LPS-exposed (hatched bars) females (light) and males (dark grey), with total number of samples in the bottom of each bar. All image analysis was completed using a custom MATLAB script and quantified the following characteristics (all units in pixels) E) Number of cells, F) Cell radius, G) Soma Area, H) Number of primary branches, I) Branch points or secondary branches, J) Total branch length. Effect size statistics were run and can be seen in Table 3, large = ###, moderate = ##, small = #.

**Table 2.** Effect size (Cohen’s ds) of microglia characteristics at P7.

|  | FSAL VS FLPS | MSAL VS MLPS | FSAL VS MSAL |
| --- | --- | --- | --- |
| Cell number | Null (0.02) | Small (0.26) | Large (2.94) |
| Cell radius | Large (0.80) | Large (1.07) | Large (2.45) |
| Soma area | Small (0.41) | Large (0.89) | Large (0.96) |
| Number of primary branches | Large (0.91) | Large (1.49) | Large (3.15) |
| Branch points | Large (1.46) | Large (1.12) | Large (3.12) |
| Total branch length | Moderate (0.50) | Large (1.18) | Large (3.40) |

**Table 3.**
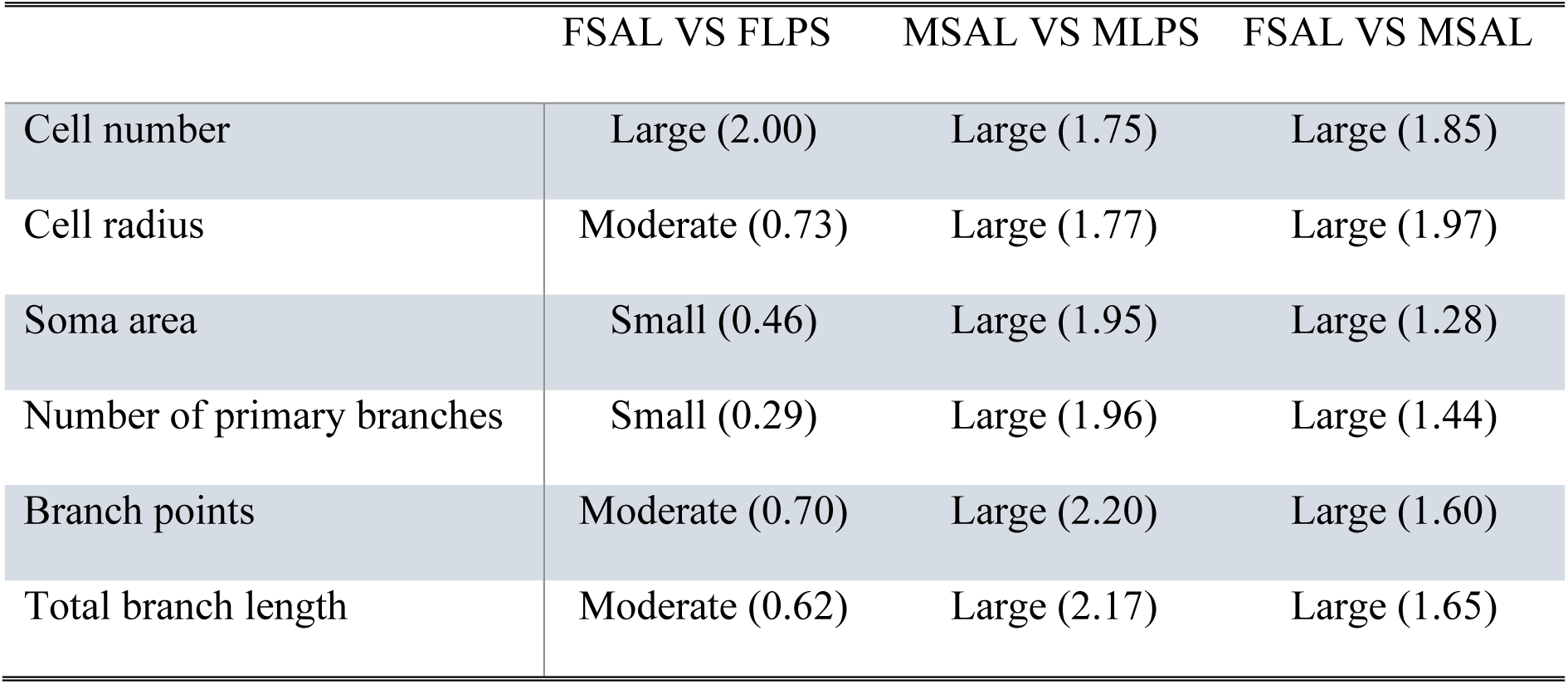
Effect size (Cohen’s d_s_) of microglial characteristics at P90.

|  | FSAL VS FLPS | MSAL VS MLPS | FSAL VS MSAL |
| --- | --- | --- | --- |
| Cell number | Large (2.00) | Large (1.75) | Large (1.85) |
| Cell radius | Moderate (0.73) | Large (1.77) | Large (1.97) |
| Soma area | Small (0.46) | Large (1.95) | Large (1.28) |
| Number of primary branches | Small (0.29) | Large (1.96) | Large (1.44) |
| Branch points | Moderate (0.70) | Large (2.20) | Large (1.60) |
| Total branch length | Moderate (0.62) | Large (2.17) | Large (1.65) |

At P7, LPS-exposed females showed a decrease in cell radius, number of primary branches and branch points of Iba1-positive cells (Fig. 5F, H, I). Cohen’s d_s_ determined these to all be large effects, shown in Table 2. A moderate and small effect of LPS-treatment was seen in total branch length (Fig. 5J) and soma area respectively (Fig. 5G), with LPS-exposed females decreasing compared to saline-exposed females. There was no effect of LPS exposure on cell number in any group (Fig. 5E). In LPS-exposed males there was a large effect in all morphological characteristics (Fig. 5F-J) beside cell number (Fig. 5E), which was a small effect (Table 2). Similar to females, microglia from LPS-exposed males exhibited decreases across all characteristics compare to saline-exposed males. Interestingly, there was a large effect in all morphological characteristics measured between the saline-treated males and females, highlighting that under normal conditions male and female microglia in the DMX show sex-specific morphology from this early age.

Neonatal LPS-exposure had an effect on adult female microglia within the DMX, with effects in all microglia characteristics observed (Table 3). LPS-exposure had a large effect on cell number in adult females, with more cells observed compared to saline-treatment (Fig. 6E). Soma area in adult females exposed to neonatal LPS also increased compared to saline-exposed females, however this was a small effect. Cell radius (Fig. 6F), branch points (Fig. 6I) and total branch length (Fig. 6J) showed a moderate decrease in adult females following neonatal LPS. The number of primary branches (Fig. 6H) also decreased in LPS-exposed females, with a small effect. In adult males there was a large effect of neonatal exposure to LPS on all measured morphological characteristics (Fig. 6E-J; Table 3). However, most impacts were observed to occur in the opposite direction to adult females. Adult males exposed to neonatal LPS showed a decrease in cell number and soma area, with accompanying increases in cell radius, number of primary branches, branch points and total branch length compared to their saline-treated counterparts. Similar to P7, there was a large effect in all characteristics between the saline-exposed males and females, providing further evidence for the sex-specificity of microglia within the DMX.

## 4. Discussion

The ability of early life events to have life-long impacts is well documented (Dinel et al., 2014), however, to date, most studies have focussed on inflammation in peripheral structures and changing behaviours in adulthood (Ellis et al., 2005). Although we know systemic neonatal inflammation modulates neuroinflammation within the brain (Wang et al., 2013), little is known about the effects of early life inflammation on the autonomic control centres within the brainstem critical to the proper physiological function of visceral organs. Here we examined whether systemic neonatal exposure to LPS affected neuroinflammation in the brainstem, and whether these changes persisted into adulthood. These data represent one of the first comprehensive explorations of a varied suite of inflammatory mediators, extending past the classic markers of inflammation, within the region of the brainstem containing a number of critical autonomic reflex centres. We found alternate responses of IL-6 expression between peripheral and CNS tissues, with the brainstem demonstrating greater sex differences than the spleen. Moreover, all inflammatory mediators showed strong sex-specific differences in the brainstem, even in the early postnatal period. LPS-exposed neonatal males showed a decrease in inflammatory mediators, whereas LPS-exposed females showed an increase. In adulthood, not all inflammatory mediators were altered by neonatal LPS-exposure, however, mediators that were altered showed contrasting results to the neonatal response, with decreases in LPS-exposed adult females and increases LPS-exposed adult males. Even more strikingly, within the brainstem, there were baseline sex differences in the majority of inflammatory mediators, indicating that naïve males and females express dramatically different levels of a number of inflammatory mediators. We also observed large effects of both neonatal LPS-exposure and sex on several morphological parameters of microglia in both acute and long-term responses, providing evidence for a potential pathway for modulation of neuroinflammation within the brainstem. Together, this data indicates that a neonatal peripheral inflammatory event can have ongoing impacts on the critical physiological control centres in the brainstem which could lead to life-long dysfunction.

### 4.1 The opposing IL-6 response of peripheral and central immune systems to neonatal LPS

The peripheral response to LPS has been well characterised (Remick et al., 2000), with the pro-inflammatory cytokine IL-6 known to be acutely upregulated following exposure at any age (Tanaka et al., 2014). To date most studies have focussed on peripheral IL-6 levels, therefore we examined the expression in the spleen in order to compare peripheral and central responses to early life LPS exposure in the same laboratory model. In males, the acute response to a neonatal stimulus in this study displayed a significant increase in spleen and a decrease in the brainstem IL-6 expression following LPS exposure. In contrast, we observed no significant differences in LPS-exposed neonatal females. It has been shown that the murine brain immune response to LPS occurs slower than in the periphery (Erickson and Banks, 2011), however, the majority of these studies have only examined adult male mice. In neonates, the immune response is tightly regulated and complex, and can produce a strong reaction in response to a stimulus. There is, however, debate as to whether this response is helpful or harmful, with initial hyper-inflammation been shown to be protective against systemic sepsis but linked to long-term CNS dysfunction (Herz et al., 2022), clearly indicating a tissue-specific role for elevated immune mediators. As the immune system develops, the peripheral and central immune systems maintain a strong bi-directional relationship, however, the intricacies of this cross-talk remain elusive (Tian et al., 2012). Ultimately, both LPS-exposed males and females showed increased expression of IL-6, indicating the long-term outcomes of IL-6 expression are similar between the sexes. In addition, in the adult brainstem, there were no significant differences in IL-6 expression in either sex. Although IL-6 is known to heavily regulate the initial phase of LPS stimulation, the long-term effects of LPS on the expression of IL-6 within the brainstem appears negligible (Wang et al., 2013).

### 4.2 Sex-specificity of inflammation in the brainstem

A key finding of this study is the unexpected baseline sex differences observed in brainstem inflammatory mediators of control animals, even during the neonatal period. In saline-exposed animals, significant sex differences were observed in IL-6, NLRP3, IL-1β, IL-10, BDNF and IFNγ expression at one week postnatal, with males having greater expression than females. The reason for sexual dimorphism has classically been attributed to sex hormones in adult models (Shepherd et al., 2020), with inflammatory mediator expression linked to sex-hormone receptors, for example IL-6 regulation by oestrogen (Miller et al., 2010). Considering P7 is well before the sexual maturity of rats (approximately 6 weeks of age) (Sengupta, 2013), the action of sex-hormones should be limited, and therefore the changes seen at this early age arise via an alternate mechanism. BDNF, one of the most well characterised neurotrophic factors, is essential to the development of the neuronal environment, especially relating to neuronal plasticity, neurotransmitter modulation and overall health to maintain normal brain function (Colucci-D’Amato et al., 2020). BDNF is also involved in many inflammatory pathways and has be shown to be sexually dimorphic pre-sexual maturity (Sardar et al., 2021). However, Sardar et al. not only showed sex-related differences but also differing expression of BDNF based on brain region, exact age and left vs right hemispheres of the brain. Given the large sex differences between inflammatory mediators at baseline, this serves to further highlight how disruption of expression during this critical developmental window can forever alter their expression in a sex-specific manner. These changes can then go on to cause life-long dysfunction, in a tissue- and sex-dependent manner, further emphasizing the need to examine all CNS regions, including those associated with autonomic control of visceral organs.

In addition to baseline sex differences in saline-treated animals, we also saw opposing responses between the sexes in response to early life LPS. At P7, LPS-exposed females showed a significant increase in NLRP3 and IL-1β, whilst males decreased their expression of these mediators following LPS. NLRP3 has two steps to complete its formation; priming and activation. The priming step is responsible for the upregulation of NLRP3 gene expression in response to stimulation via LPS (Blevins et al., 2022). A second stimulus then initiates the activation step, and once active, NLRP3 activates caspase-1 which begins the release of IL-1β (Kelley et al., 2019). Given the upregulation of both NLRP3 and IL-1β in this study it indicates that this inflammasome pathway is more active in females, and downregulated in males. The NLRP3 inflammasome pathway plays a key role in innate immunity and providing a robust inflammatory defence to pathogens, implicating its activity as a key driver of neuroinflammation and associated damage (Song et al., 2017). Although early life LPS exposure strongly altered the NLRP3 pathway during the neonatal period, minimal changes were observed in adulthood, suggesting the long-term impact may not be as important as the acute responses, however, the effect of such strong modulation during this critical window of development requires further exploration. In addition, LPS-exposed neonatal males displayed a significant decrease in all inflammatory mediators examined in this study. Males are known to have poorer outcomes and increased mortality from neonatal infections, and this is thought to be derived from high levels of testosterone that suppress the immune system (Aghai et al., 2020, Subedi et al., 2022, Townsel et al., 2017). Therefore, this suppression of inflammatory mediators within the brainstem could be implicated in autonomic dysfunction and subsequent complications of infection.

In adulthood, females exposed to neonatal LPS showed no changes in pro-inflammatory mediators, however there was a significant decrease in IL-1β and IFNγ. There was also a significant decrease in the anti-inflammatory mediators, BDNF and IL-10, which are responsible for reducing inflammation arising from future insults and are especially important for optimal functioning and maintenance of neuronal networks (Miranda et al., 2019, Nenov et al., 2019). In contrast, adult males exposed to early life LPS displayed a significant increase in NLRP3 expression and, as stated above, an increase in gene expression indicates NLRP3 is in the primed stage, awaiting activation (Blevins et al., 2022). In the event of a second exposure to an inflammatory agent, this would suggest that males would have a larger, faster inflammatory response, as they are now in a ‘primed’ state. In contrast, given females have lowered levels of both anti- and pro-inflammatory mediators, would likely show an attenuated, but potentially prolonged, response to exposure to a second inflammatory agent as they would not be positioned to mount an appropriate inflammatory response to clear the stimulus. This is seen in humans with men being more susceptible to an acute infection and more likely to produce an acute response, whereas women show more persistent immune responses with greater risk of chronic inflammatory conditions (Chamekh and Casimir, 2019). Although little is known about the functional consequences of altered inflammatory mediators in autonomic brainstem networks (Hedley et al., 2022), it is clear that the sexually dimorphic responses observed at both P7 and P90 would likely have serious implications on visceral physiology for both males and females.

### 4.3 Microglia are a potential contributory mechanism for ongoing neuroinflammation

Microglia are essential to synaptic formation and pruning in the developmental stage (Lenz and Nelson, 2018) and disruptions to this function have been linked to neurodevelopmental disorders (Nelson and Lenz, 2017) and neuronal cell death (Ueno et al., 2013) Therefore, we applied a custom MATLAB script (Abdolhoseini et al., 2019, Abdolhoseini et al., 2016) to Iba1+ve microglia in order to examine microglial morphology. Our data strongly indicate that microglia are contributing to both immediate and long-term modulation resulting from neonatal LPS exposure. At P7, effects were seen across all morphological characteristics, with both male and female microglia displayed similar modulations, although LPS had a larger effect on male microglia. Overall, in LPS-exposed animals, microglia showed acute responses of decreases in branching and cell radius compared to their saline counterparts. Using traditional classification methods, the morphology of these cells would be classed between activated and amoeboid, however, our automated approach allowed for a more detailed assessment of their morphology. In our sample, microglia have retracted their branches but are not in full amoeboid morphology, which would be indicated by an increase in soma size (Colonna and Butovsky, 2017) that was not observed in our analysis. It has been previously described that neonatal LPS delays the developmental change from amoeboid to ramified microglia morphology over the first week of life (Cardoso et al., 2015). Our data further supports this finding, and also adds to the growing body of literature regarding the complexity of sex-specific responses, even in these early postnatal ages.

In adulthood, we observed modulation of microglial morphology following neonatal LPS exposure that presented in a sexually dimorphic manner. In adult males, LPS-exposure had a large effect on all measured microglial characteristics, with a decrease in cell number and soma area, coupled with the increase in cell radius and branching indicates a larger network of branching. This morphological state is traditionally referred to as a primed, or hyper-ramified, state (Lecours et al., 2018). Microglia in this primed state are more susceptible to a milder second stimulus or will have an exaggerated reaction to a stimulus of similar magnitude to the first (Lima et al., 2022). This type of exaggerated response to a second hit can result in further damage to the CNS leading to greater dysfunction of homeostatic control and synaptic rigidity (Fernández-Arjona et al., 2022, Pires et al., 2020), which aligns with the modulation of the expression of inflammatory mediators we observed in the adult male brainstem following early life LPS exposure. In contrast to males, adult females demonstrated increased numbers of microglia within key autonomic nuclei in the brainstem following early life LPS exposure, with a small to moderate effect on all other measured features. Overpopulation of microglia can lead to excessive production of inflammatory mediators in response to an stimulus, which can ultimately have neurodegenerative effects (Bachiller et al., 2018). Surprisingly, this counters our finding of decreased inflammatory mediators in the adult female brainstem, suggesting the role of microglia may be less tightly linked to the inflammatory profile of the brainstem in adult females than males.

Additionally, our data revealed a large effect of sex on all microglial characteristics measured at both ages examined, with males and females exposed to saline demonstrating different microglial morphology. Saline exposed female microglia had a larger cell radius and more branching than saline-exposed male microglia, whereas the male DMX had more cells with larger soma area than the female DMX. Together, this suggests that although males have more microglia, they are closer to the ameboid, or activated, end of the neuroanatomical spectrum than females (Leyh et al., 2021). This would suggest male microglia are more likely to respond to stimulation, once again indicating males are closer to threshold at rest than females. In addition, adult microglia females typically showed a neuroprotective phenotype by expressing anti-inflammatory markers, and when transplanted into a male brain, the female microglia retained the female morphology, indicating that their morphology was independent of the hormonal environment (Villa et al., 2018). The LPS-induced modulation of female microglia observed in this study could be linked to the decreased expression of anti-inflammatory markers in adult females exposed to early life LPS. However, further investigation into the specificity of the altered microglial activity is warranted to further explore their role in the life-long implications of early life LPS exposure.

## 5. Conclusions

This study highlights the importance of the peripheral to central immune pathway and how disruption of this pathway during development can have lifelong consequences. Baseline differences in inflammatory mediators and microglial morphology between the sexes, as well as differing responses to early life LPS exposure, clearly indicate a need for studying males and females equally at all ages. Overall, neonatal LPS significantly increases the inflammatory profile of the brainstem in a sex-specific manner. In the short term, this can lead to substantial neurodevelopmental challenges and in the long-term, a predisposition to further damage. Given the role of the brainstem in providing innervation to viscera, enhanced neuroinflammation within this region could have wide-reaching, sexually dimorphic implications for many physiological functions, including airway patency and gastric motility (Fontán et al., 2000, Zhou et al., 2008).

## Authors’ contributions

All authors conceived and designed the study. KEH performed all experiments and analysis, prepared all figures and wrote the manuscript. AC ran the laboratory model and aided in formulating the statistical parameters. MAT reviewed all data and proofread the manuscript. RJC, DMH and JCH proofread the manuscript and provided mentoring to the ECR authors. All authors read and approved the final manuscript.

