## Supplemental figures 1-4 for "Autonomic regions of the brainstem show a sex-specific inflammatory response to systemic neonatal lipopolysaccharide"

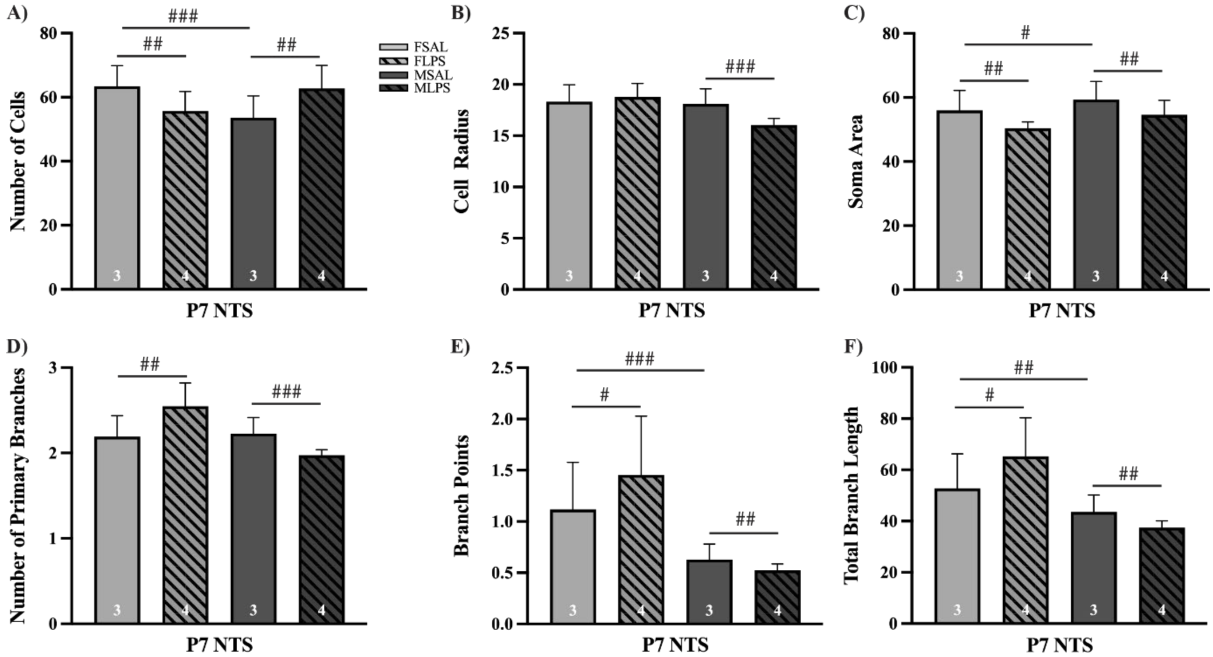


***Supplementary figure 1. Immunofluorescence of microglia within the solitary tract of the nucleus at P7.*** Bar graphs showing microglia morphological characteristics (mean ± SEM) for Saline-exposed (open bars) and LPS-exposed (hatched bars) females (light) and males (dark grey), with total number of samples in the bottom of each bar. All image analysis was completed using a custom MATLAB script and quantified the following characteristics (all units in pixels) A) Number of cells, B) Cell radius, C) Soma Area, D) Number of primary branches, E) Branch points or secondary branches, F) Total branch length. Effect size statistics were run and can be seen in Table 3, large = ###, moderate = ##, small = #.

***Supplementary table 1. Effect size (Cohen's $d_s$) of microglia characteristics in NTS at P7***

| | FSAL VS FLPS | MSAL VS MLPS | FSAL VS MSAL |
|---|---|---|---|
| Cell number | Moderate (0.65) | Moderate (0.67) | Large (0.84) |
| Cell radius | Null (0.17) | Large (1.07) | Null (0.09) |
| Soma area | Moderate (0.74) | Moderate (0.50) | Small (0.33) |
| Number of primary branches | Moderate (0.70) | Large (1.10) | Null (0.09) |
| Branch points | Small (0.32) | Moderate (0.55) | Large (0.82) |
| Total branch length | Small (0.44) | Moderate (0.73) | Moderate (0.50) |

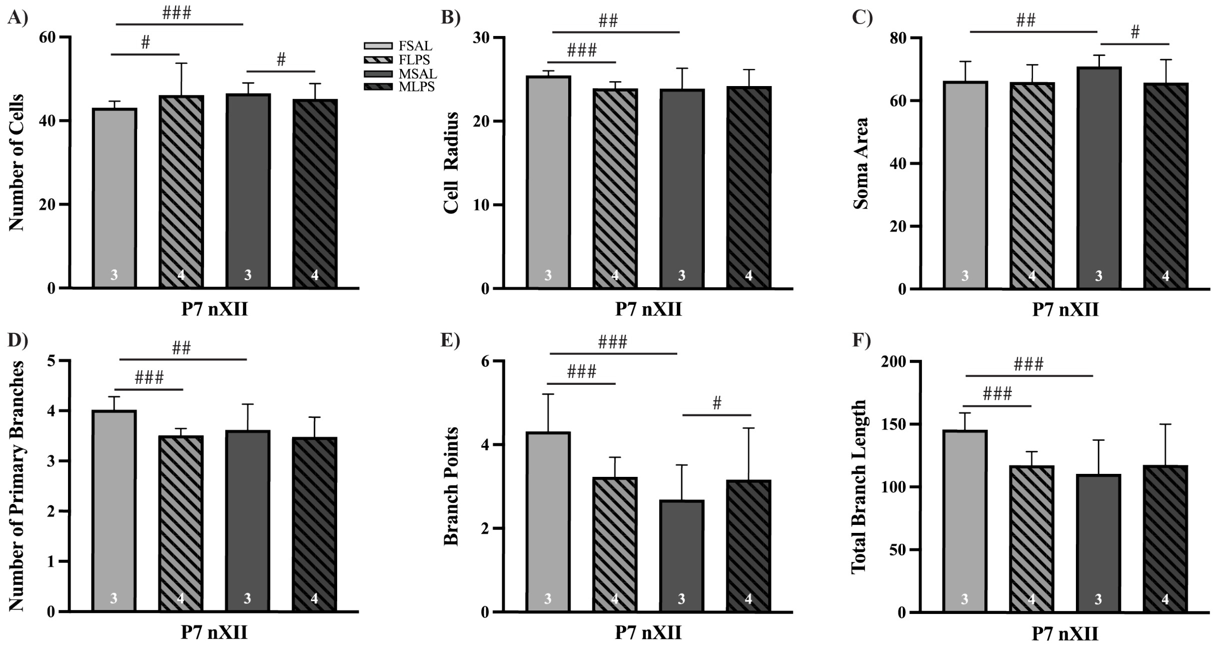


***Supplementary figure 2. Immunofluorescence of microglia within the hypoglossal nucleus at P7.*** Bar graphs showing microglia morphological characteristics (mean ± SEM) for Saline-exposed (open bars) and LPS-exposed (hatched bars) females (light) and males (dark grey), with total number of samples in the bottom of each bar. All image analysis was completed using a custom MATLAB script and quantified the following characteristics (all units in pixels) A) Number of cells, B) Cell radius, C) Soma Area, D) Number of primary branches, E) Branch points or secondary branches, F) Total branch length. Effect size statistics were run and can be seen in Table 3, large = ###, moderate = ##, small = #.

***Supplementary table 2. Effect size (Cohen's $d_s$) of microglia characteristics in nXII at P7***

| | FSAL VS FLPS | MSAL VS MLPS | FSAL VS MSAL |
|---|---|---|---|
| Cell number | Small (0.25) | Small (0.21) | Large (0.95) |
| Cell radius | Large (1.11) | Null (0.08) | Moderate (0.50) |
| Soma area | Null (0.03) | Small (0.42) | Moderate (0.52) |
| Number of primary branches | Large (1.40) | Null (0.17) | Moderate (0.56) |
| Branch points | Large (0.88) | Small (0.22) | Large (1.08) |
| Total branch length | Large (1.25) | Null (0.12) | Large (0.94) |

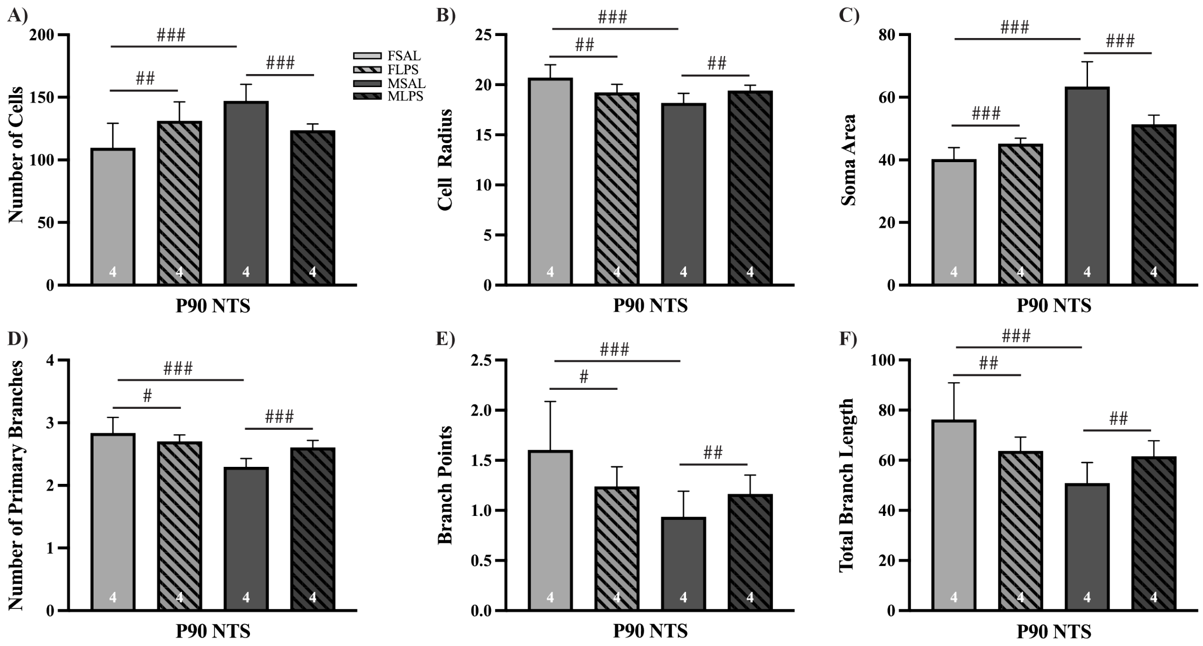


***Supplementary figure 3. Immunofluorescence of microglia within the solitary tract of the nucleus at P90.*** Bar graphs showing microglia morphological characteristics (mean ± SEM) for Saline-exposed (open bars) and LPS-exposed (hatched bars) females (light) and males (dark grey), with total number of samples in the bottom of each bar. All image analysis was completed using a custom MATLAB script and quantified the following characteristics (all units in pixels) A) Number of cells, B) Cell radius, C) Soma Area, D) Number of primary branches, E) Branch points or secondary branches, F) Total branch length. Effect size statistics were run and can be seen in Table 3, large = ###, moderate = ##, small = #.

***Supplementary table 3. Effect size (Cohen's $d_s$) of microglia characteristics in NTS at P90***

| | FSAL VS FLPS | MSAL VS MLPS | FSAL VS MSAL |
|---|---|---|---|
| Cell number | Moderate (0.61) | Large (1.56) | `Large (1.11) |
| Cell radius | Moderate (0.69) | Moderate (0.77) | Large (1.11) |
| Soma area | Large (0.84) | Large (0.99) | Large (1.84) |
| Number of primary branches | Small (0.34) | Large (1.20) | Large (1.32) |
| Branch points | Small (0.49) | Moderate (0.50) | Large (0.85) |
| Total branch length | Moderate (0.56) | Moderate (0.73) | Large (1.06) |

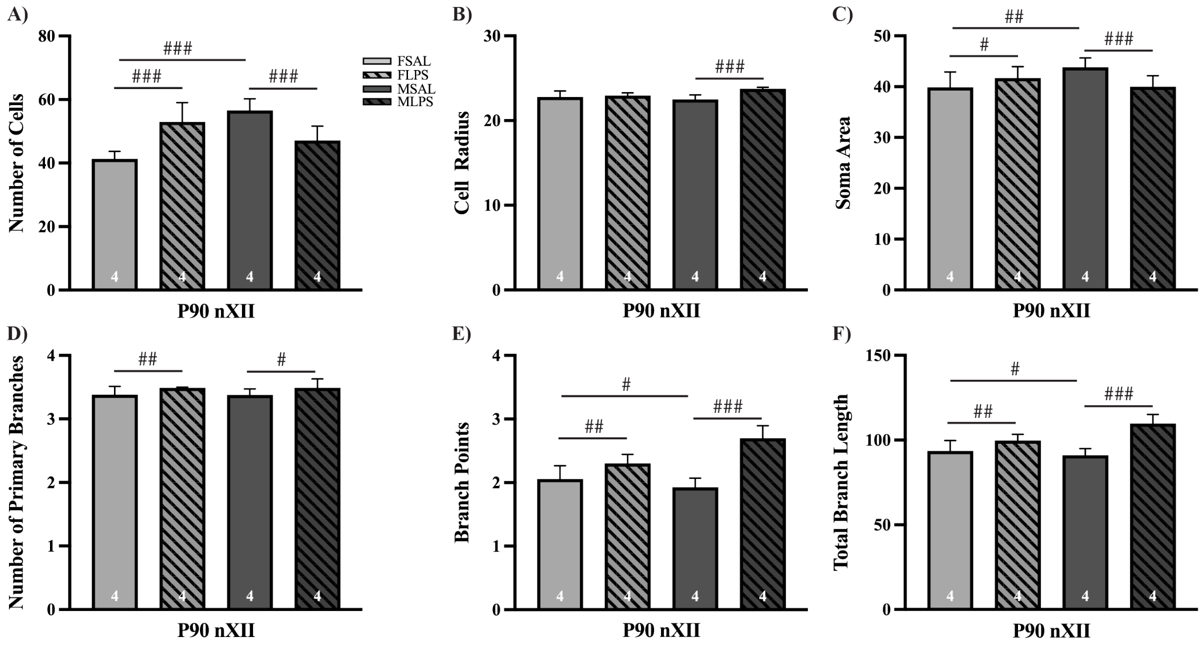


***Supplementary figure 4. Immunofluorescence of microglia within the hypoglossal nucleus at P90.*** Bar graphs showing microglia morphological characteristics (mean ± SEM) for Saline-exposed (open bars) and LPS-exposed (hatched bars) females (light) and males (dark grey), with total number of samples in the bottom of each bar. All image analysis was completed using a custom MATLAB script and quantified the following characteristics (all units in pixels) A) Number of cells, B) Cell radius, C) Soma Area, D) Number of primary branches, E) Branch points or secondary branches, F) Total branch length. Effect size statistics were run and can be seen in Table 3, large = ###, moderate = ##, small = #.

***Supplementary table 4. Effect size (Cohen's $d_s$) of microglia characteristics in nXII at P90***

| | FSAL VS FLPS | MSAL VS MLPS | FSAL VS MSAL |
|---|---|---|---|
| Cell number | Large (1.23) | Large (1.12) | Large (2.40) |
| Cell radius | Null (0.16) | Large (1.45) | Null (0.19) |
| Soma area | Small (0.33) | Large (0.92) | Moderate (0.77) |
| Number of primary branches | Moderate (0.54) | Small (0.44) | Null (0.01) |
| Branch points | Moderate (0.65) | Large (2.15) | Small (0.35) |
| Total branch length | Moderate (0.58) | Large (1.90) | Small (0.24) |
